# Ligand-independent modulation of GIPR signaling by splice variants

**DOI:** 10.1101/2022.01.24.477496

**Authors:** Kaini Hang, Lijun Shao, Qingtong Zhou, Fenghui Zhao, Antao Dai, Xiaoqing Cai, Raymond C. Stevens, Dehua Yang, Ming-Wei Wang

## Abstract

Glucose-dependent insulinotropic polypeptide receptor (GIPR) is a potential drug target for metabolic disorders. It works with glucagon-like peptide-1 receptor (GLP-1R) and glucagon receptor (GCGR) in humans to maintain glucose homeostasis. Unlike the other two receptors, GIPR has at least 7 reported (EMBL-EBI, 2022; NCBI, 2022a, 2022b) splice variants (SVs) with previously undefined functions. To explore their roles in endogenous peptide mediated GIPR signaling, we investigated the outcome of co-expressing each of the four SVs in question with GIPR in terms of ligand binding, cAMP accumulation, G_s_ activation, β-arrestin recruitment and cell surface localization. The effects of these SVs on intracellular cAMP responses modulated by receptor activity-modifying proteins (RAMPs) were also studied. It was found that while SVs of GIPR neither bound to the hormone nor affected its signal transduction *per se*, they differentially regulated GIPR-mediated cAMP and β-arrestin responses. Specifically, SV1 and SV4 were preferable to G_s_ signaling, SV3 was biased towards β-arrestin recruitment, whereas SV2 was inactive on both pathways. In the presence of RAMPs, only SV1 and SV4 synergized the repressive action of RAMP3 on GIP-elicited cAMP production. The results suggest a new form of signal bias that is constitutive and ligand-independent, thereby expanding our knowledge of biased signaling beyond pharmacological manipulation (*i*.*e*., ligand specific) as well as constitutive and ligand-dependent (*e*.*g*., SV1 of the growth hormone-releasing hormone receptor).

## Introduction

G protein-coupled receptors (GPCRs) represent the largest family of membrane proteins that are universally expressed in human tissues (Pavlos & Friedman, 2017). They can recognize a diverse range of extracellular ligands and transduce signals to intracellular coupling partners, thereby governing crucial physiological functions (Strange, 2008). GPCR-mediated signaling and pharmacological activities could be profoundly affected by alternative splicing, leading to functional diversity (Furness, Wootten, Christopoulos, & Sexton, 2012; Marti-Solano et al., 2020).

Splice variants (SVs) have been observed in many GPCRs, in which sequence variations may include N terminus truncation or/and substitution, C terminus truncation or/and substitution, intracellular/extracellular loop differences, severe truncation leading to variants with less than 7 transmembrane domains (TMDs) or soluble variants (Markovic & Challiss, 2009). In general, N terminus variations impair ligand binding properties, such as corticotropin releasing hormone receptor 1 (CRH1R) and 2 (CRHR2) (Evans & Seasholtz, 2009), calcitonin receptor (CTR) (Nag, Sultana, Kato, & Hirose, 2007) and parathyroid hormone 1 receptor (PTH1R) (Joun et al., 1997), while C terminus variations show altered signaling or protein interactions, such as metabotropic glutamate receptors (mGluRs) (Cai, Schools, & Kimelberg, 2000), μ-opioid receptor (MOR) (Lu et al., 2015) and 5-hydroxytryptamine (5-HT) receptors (Coupar, Desmond, & Irving, 2007). Intracellular loop (ICL) differences affect G protein coupling preference and extracellular loop (ECL) differences alter ligand specificity and binding kinetics, as illustrated by pituitary adenylate cyclase-activating polypeptide (PACAP) type 1 receptor (PAC1R) (McCulloch et al., 2001) and D3 dopamine receptor (D3R) (Karpa, Lin, Kabbani, & Levenson, 2000; Richtand, 2006), respectively. Variants with less than 7 TMDs caused by severe N terminus truncation exhibit negative effect on wild-type (WT) receptor signaling (Markovic & Challiss, 2009).

Glucose-dependent insulinotropic polypeptide receptor (GIPR) belongs to class B1 subfamily of GPCRs and is present in pancreatic cells, adipose tissues and osteoblasts. Upon GIP stimulation, it regulates insulin secretion, fat accumulation and bone formation by increasing intracellular adenosine 3,5-cyclic monophosphate (cAMP) levels (Campbell, 2021; Yabe & Seino, 2011; Yue & Lam, 2019). GIPR is reported to have a truncated SV showing a dominant negative effect on the translocation of WT GIPR from the endoplasmic reticulum (ER) to the cell surface, leading to a decreased activity (Harada et al., 2008). However, the functionalities of GIPR SVs remain to be defined.

In this study, we constructed and expressed four representative GIPR SVs to elaborate their biological properties on GIPR-mediated cAMP accumulation and β-arrestin recruitment. They were selected based on expression levels and splicing modes: SV1, SV2, SV3 and SV4 with residue lengths of 419, 430, 405 and 265, respectively (Figure 1). SV1 has a truncated C terminus and a 20 amino acid substitution. SV2 lacks the sequence of residues 58-93 at the N terminus. SV3 has a replaced N terminus of residues 1-93. SV4 only has 3 TMDs.

**Figure 1.**
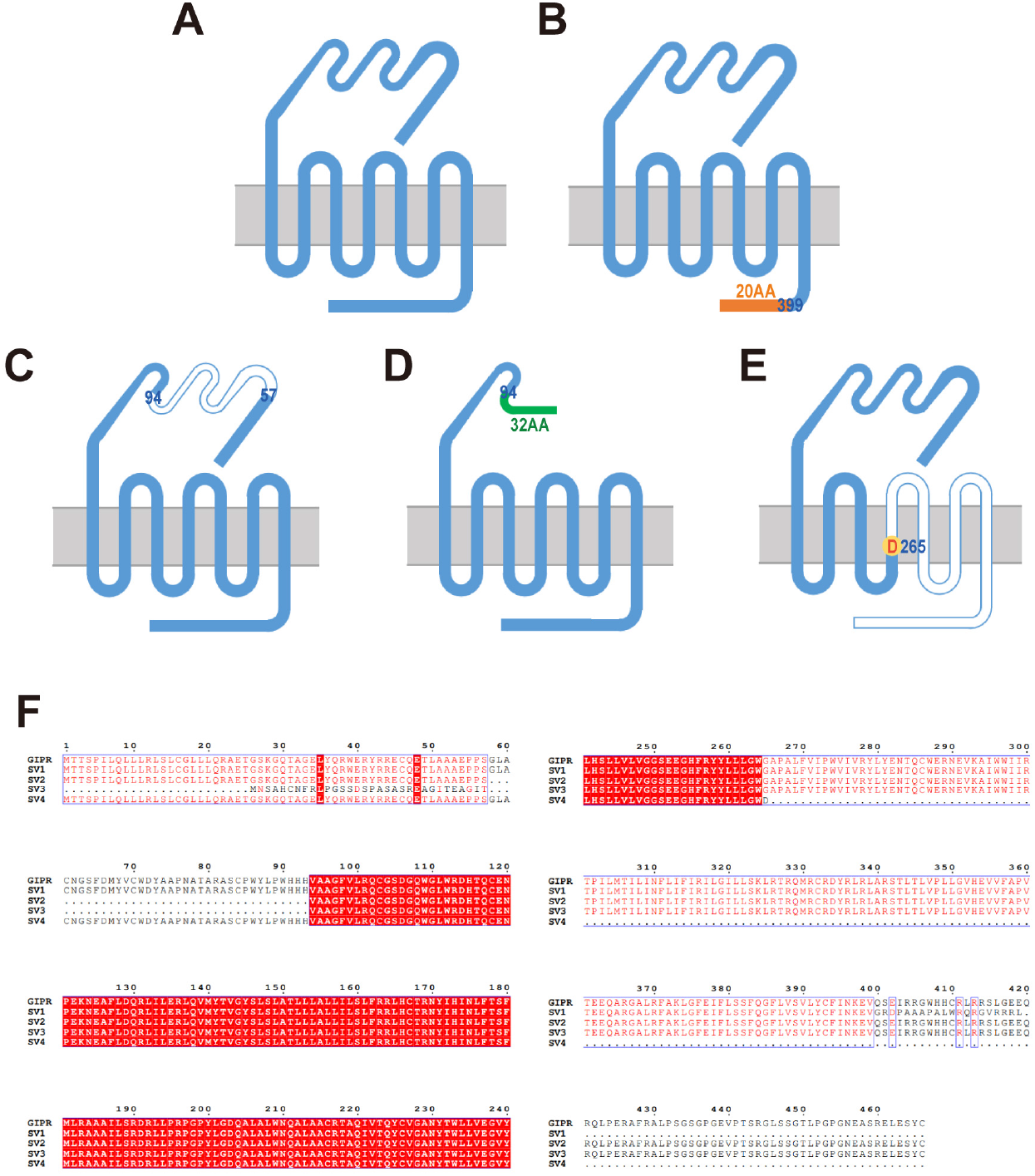
Schematic diagram of GIPR and its splice variant (SV) constructs. **A**, Construct of the wild-type (WT) GIPR. **B**, Construct of SV1. Residues 400-466 are replaced by a 20-amino acid sequence (GRDPAAAPALWRQRGVRRRL). **C**, Construct of SV2. Residues 58-93 are missing compared to that of WT. **D**, Construct of SV3. Residues 1-93 are replaced by a 32-amino acid sequence (MNSAHCNFRLPGSSDSPASASREAGITEAGIT). **E**, Construct of SV4. Residues 266-466 are missing and G265 is replaced by aspartic acid (D). **F**, Sequence alignment of GIPR and the four SVs.

## Results

### Splice variants neither bind nor affect GIP-induced cAMP response

We first expressed GIPR and four SVs separately in HEK293T cells to investigate their ability to bind GIP_1-42_ and elicit cAMP accumulation and β-arrestin recruitment. Figure 2 shows that none of the SVs displayed any ligand-binding and signaling properties, whereas the WT receptor was highly active in each parameter analyzed.

**Figure 2.**
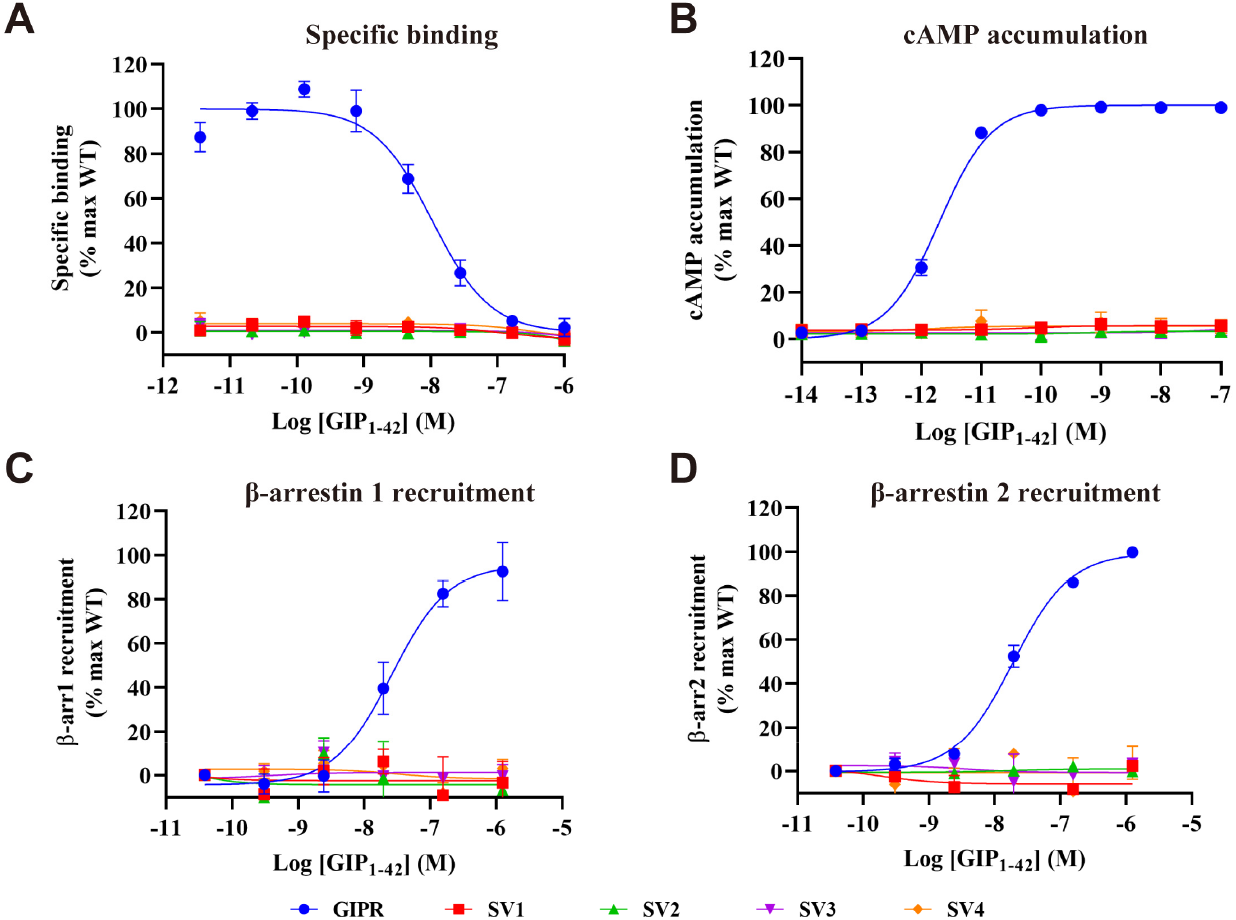
Ligand-binding and signaling profiles of GIPR and its splice variants (SVs). **A**, Competitive inhibition of ^125^I-GIP_1-42_ binding to GIPR and SVs by unlabeled GIP_1-42_. Binding affinity is quantified by reduction of radioactivity (counts per minute, CPM). **B**, Concentration-response curves of cAMP accumulation elicited by GIP_1-42_ at GIPR and SVs. **C and D**, β-arrestins 1 (β-arr1) and 2 (β-arr2) recruitment by GIPR and SVs. Concentration-response characteristics are shown as the area-under-the-curve (AUC) across the time-course response curve (0 to 10 min) for each concentration. Data shown are means ± SEM of at least three independent experiments (n=3-5) performed in quadruplicate (cAMP accumulation) or duplicate (specific binding and β-arrestin recruitment). Signals were normalized to the maximum (max) response of the wild-type (WT) GIPR and concentration-response curves were analyzed using a three-parameter logistic equation. The following figure supplements are available for figure 2: Source data 1. Ligand-binding and signaling profiles of GIPR and its splice variants (SVs).

### Splice variants differentially modulate GIPR-mediated signaling

Since SVs are usually express in cells and tissues where GIPR are present (GTEx, 2022; Harada et al., 2008), we co-transfected GIPR with each SV in order to study if they influence the signaling profile of the WT receptor. While the binding affinity of GIP_1-42_ to the cognate receptor were significantly reduced by 0.44, 0.54, 0.50 and 0.74 folds for SV1, SV2, SV3 and SV4, respectively (Figure 3A and Table 1), cAMP and β-arrestin responses at GIPR were differentially and negatively modulated. Both SV1 and SV4 decreased cAMP signaling but the effect of SV4 was nearly 5-fold stronger than that of SV1, SV2 and SV3 were inactive (Figure 3B and Table 1). Although SV1, SV3 and SV4 decreased the E_max_ values of β-arrestin 2 recruitment by 0.59, 0.49 and 0.42-folds, respectively, and SV2 remained inactive (Figure 3D and Table 1), none of them influenced on β-arrestin 1 recruitment (Figure 3C).

**Table 1.** Effects of splice variants (SVs) on ligand binding and GIPR-mediated signal transduction in HEK293T cells co-expressing GIPR and individual SVs.

|  | Ligand binding |  | cAMP accumulation |  | G<sub>s</sub> coupling |  | β-arr1 recruitment |  | β-arr2 recruitment |  |
| --- | --- | --- | --- | --- | --- | --- | --- | --- | --- | --- |
|  | pIC<sub>50</sub> ± SEM | Span (%) ± SEM | pEC<sub>50</sub> ± SEM | E<sub>max</sub> (%) ± SEM | pEC<sub>50</sub> ± SEM | E<sub>max</sub> (%) ± SEM | pEC<sub>50</sub> ± SEM | E<sub>max</sub> (%) ± SEM | pEC<sub>50</sub> ± SEM | E<sub>max</sub> (%) ± SEM |
| **WT GIPR** | 7.91 ± 0.10 | 100.00 ± 4.87 | 11.14 ± 0.02 | 100.00 ± 0.63 | 9.66 ± 0.08 | 99.79 ± 2.20 | 7.94 ± 0.26 | 99.03 ± 6.83 | 8.19 ± 0.06 | 99.78 ± 1.81 |
| **GIPR+SV1** | 8.09 ± 0.09 | 54.21 ± 2.27* | 10.92 ± 0.02* | 100.08 ± 0.50 | 8.89 ± 0.24 | 91.36 ± 6.42 | 7.29 ± 0.52 | 126.65 ± 16.04 | 8.32 ± 0.35* | 40.99 ± 4.17 |
| **GIPR+SV2** | 7.96 ± 0.11 | 46.52 ± 2.30* | 11.10 ± 0.02 | 100.79 ± 0.52 | 9.39 ± 0.19 | 102.39 ± 5.10 | 7.67 ± 0.44 | 88.87 ± 9.90 | 8.21 ± 0.06 | 96.02 ± 1.69 |
| **GIPR+SV3** | 7.82 ± 0.13 | 51.01 ± 3.26* | 11.16 ± 0.02 | 100.38 ± 0.43 | 9.67 ± 0.21 | 93.61 ± 5.31 | 7.70 ± 0.39 | 96.31 ± 8.45 | 8.22 ± 0.34* | 50.62 ± 5.12 |
| **GIPR+SV4** | 8.15 ± 0.13 | 25.77 ± 1.59* | 10.46 ± 0.02* | 100.38 ± 0.68 | 9.10 ± 0.22 | 85.77 ± 5.40 | 8.08 ± 0.60 | 96.27 ± 10.78 | 8.13 ± 0.16* | 58.17 ± 2.88 |

cAMP accumulation, G<sub>s</sub> activation and β-arrestin 1/2 recruitment assays were performed in HEK293T cells. Whole cell binding assay was performed in CHO-K1 cells. All the measures were fitted to non-linear regression three-parameter logistic curves. pEC<sub>50</sub> and pIC<sub>50</sub> are the negative logarithm of the concentration of an agonist that gives a half of the maximum response. E<sub>max</sub> and span values are the percentage (%) of the maximum response in cells expressing GIPR only. Data shown are means ± SEM of at least three independent experiments. One-way ANOVA was used to determine statistical difference (\*P < 0.0001). β-arr1, β-arrestin 1; β-arr2, β-arrestin 2; WT, wild-type.

**Figure 3.**
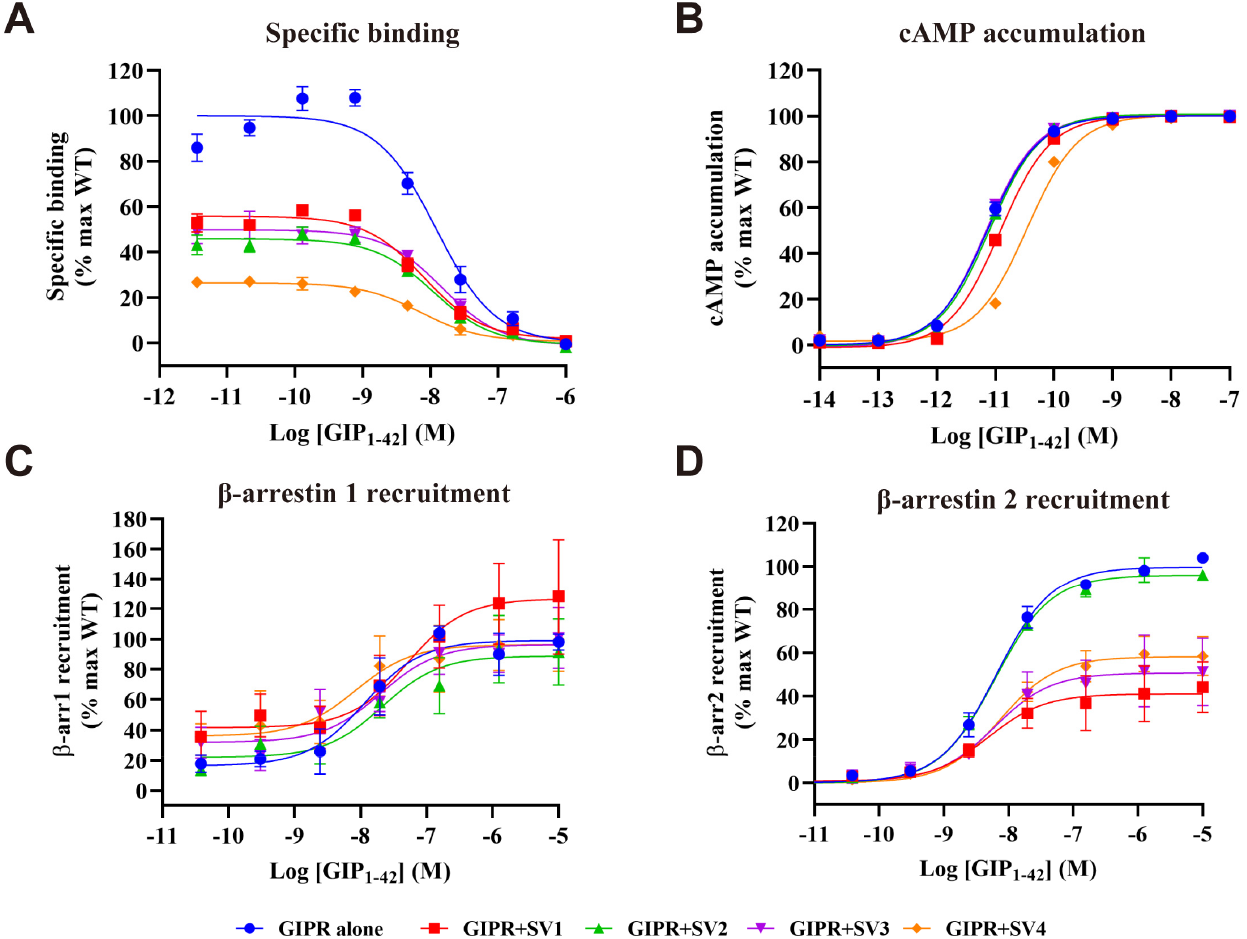
Effects of GIPR splice variants (SVs) on ligand binding and the wild-type (WT) GIPR mediated signal transduction in HEK293T cells co-expressing GIPR and individual SVs. **A**, Effects of SVs on competitive binding of ^125^I-GIP_1-42_ to GIPR. **B**, Effects of SVs on GIP_1-42_ induced cAMP accumulation at GIPR. **C and D**, Effects of SVs and GIPR on β-arrestins 1 (β-arr1) and 2 (β-arr2) recruitment by GIPR. Cells were co-transfected with GIPR and each SV at a 1:3 ratio. Data shown are means ± SEM of at least three independent experiments (n=3-5) performed in quadruplicate (cAMP accumulation) or duplicate (specific binding and β-arrestin recruitment). Signals were normalized to the maximum (max) response of GIPR and concentration-response curves were analyzed using a three-parameter logistic equation. The following figure supplements are available for figure 3: Source data 1. Effects of GIPR splice variants (SVs) on ligand binding and the wild-type (WT) GIPR mediated signal transduction in HEK293T cells co-expressing GIPR and individual SVs.

### GIPR and splice variants are co-localized on the membrane

GIPR and SVs could be localized either on the membrane or the cytoplasm of transfected HEK293T cells (Figure 4A and 4B). Figure 4C illustrates that GIPR, SV1 and SV4 were co-expressed mostly on the cell surface, whereas SV2 and SV3 only exhibited a partial membrane co-localization. Upon co-transfection with GIPR, most of SV3 were translocated to the membrane (3^rd^ panel of Figure 4C), but SV2 remained in the cytoplasm along with redistributed GIPR (2^nd^ panel of Figure 4C), consistent with the silent role of SV2 observed.

**Figure 4.**
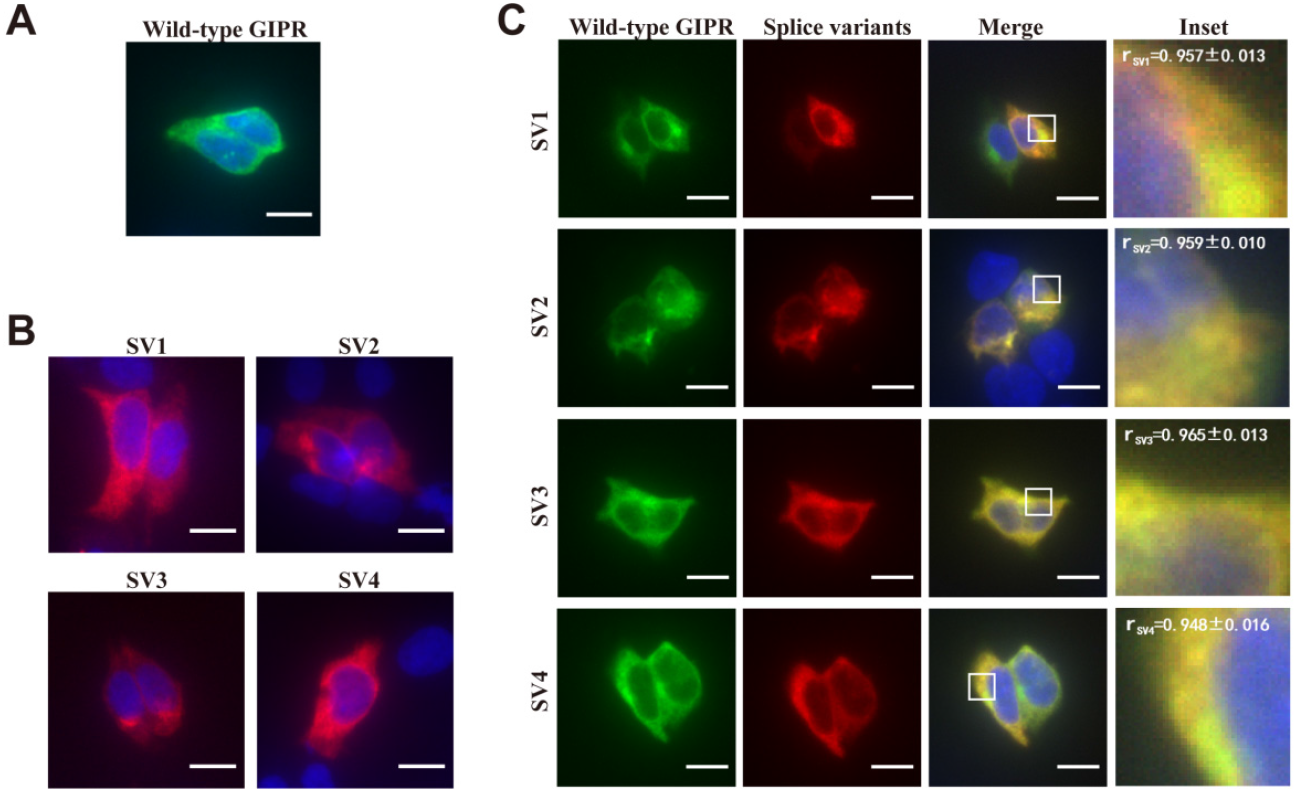
Co-localization of GIPR and its splice variants (SVs). Immunofluorescence staining of HEK293T cells transfected with GIPR-HA (**A**) or each SV-FLAG (**B**) alone. To estimate their co-localization, co-transfection of GIPR and individual SVs (**C**) was performed at a ratio of 1:3 (green, GIPR-HA; red, SV-FLAG; yellow, merge). Data show representative results from three independent experiments. Inset demonstrates the overlapping positions of GIPR and SVs in the cell surface (SV1, SV3 and SV4) or cytoplasm (SV2). Cells were observed by DeltaVision™ Ultra. Scale bar = 10 μm. The following figure supplements are available for figure 4: Source data 1. Co-localization of GIPR and its splice variants (SVs).

### Synergistic effect exerted by splice variants and RAMP3

Receptor activity-modifying protein 3 (RAMP3) was reported to reduce GIP_1-42_ induced cAMP accumulation at GIPR as opposed to RAMPs 1 and 2 that showed no effect (Harris, Mackie, Pawlak, Carvalho, & Ladds, 2021; Shao et al., 2021). After co-expression of individual SVs with GIPR and each RAMP, no alteration was noted with RAMP1 and RAMP2 (Figure 5A and 5B), but the suppression of GIPR-mediated cAMP production by RAMP3 was moderately augmented by SV1 and SV4 (with EC_50_ increased by 0.54 and 0.96-fold, respectively) (Figure 5C and Table 2).

**Table 2.**
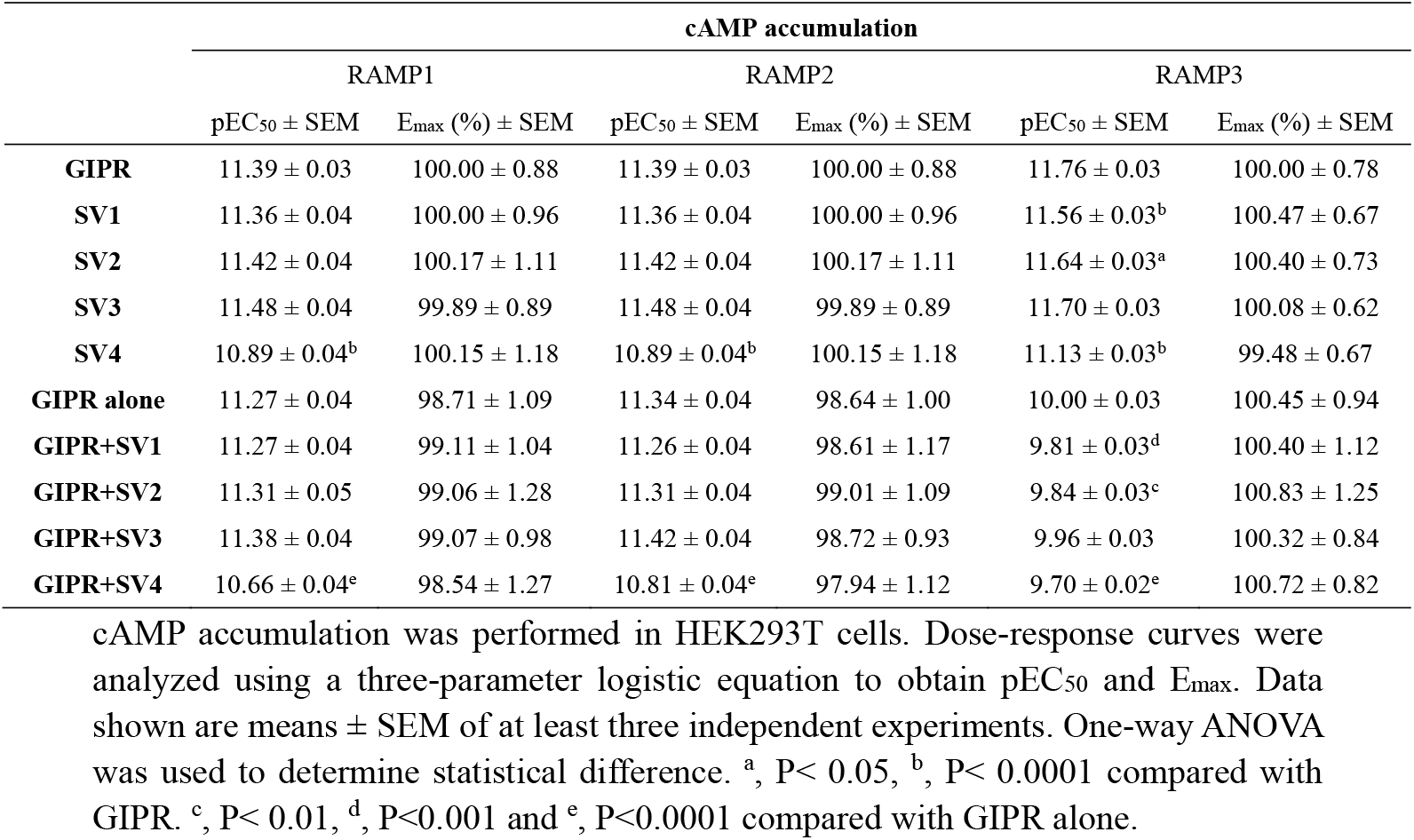
Synergistic effects of splice variants (SVs) and receptor activity-modifying 612 proteins (RAMPs) on GIPR-mediated cAMP signaling in HEK293T cells co-613 expressing GIPR.

**Figure 5.**
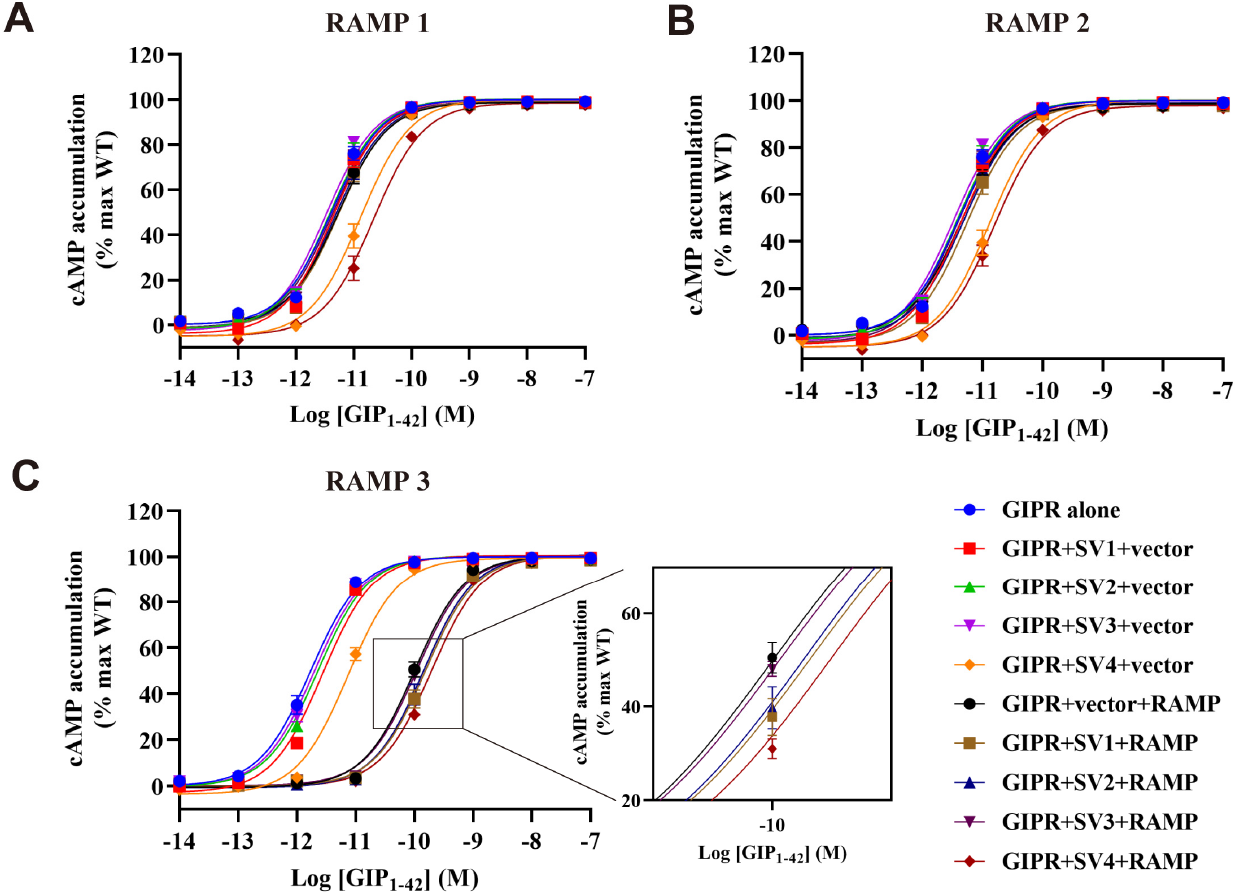
Synergistic effects of splice variants (SVs) and receptor activity-modifying proteins (RAMPs) on GIPR-mediated cAMP signaling in HEK293T cells co-expressing GIPR, each SV and RAMP1 (**A**), RAMP2 (**B**) or RAMP3 (**C**). Signals were normalized to the maximum (max) response of the wild-type (WT) GIPR and data were fitted with a non-linear regression of three-parameter logistic curve. Data shown are means ± SEM of three independent experiments performed in quadruplicate. The following figure supplements are available for figure 5: Source data 1. Effects of splice variants (SVs) and receptor activity-modifying proteins (RAMPs) on GIPR-mediated cAMP signaling in HEK293T cells co-expressing GIPR, each SV and RAMP1.

### SV1 and SV4 are repressive on G_s_ activation

We also studied the effect of SVs on G_s_ protein coupling by GIPR. G_s_ activation was assessed using a split luciferase NanoBiT G protein sensor. Individually expressing SVs showed no ability to active G_s_ (Figure 6A), consistent with their lack of cAMP signaling (Figure 2B). SV1 and SV4 markedly impaired G_s_ coupling with 4.89-and 2.68-fold increased EC_50_ values, respectively (Figure 6B and Table 1). Although the P values of pEC_50_ for SV1 and SV4 were greater than 0.05 probably due to inherent assay variations, the difference between their EC_50_ and that of GIPR alone was statistically significant (Figure 6C and D), thus in line with their repressive action on cAMP accumulation (Figure 3B).

**Figure 6.**
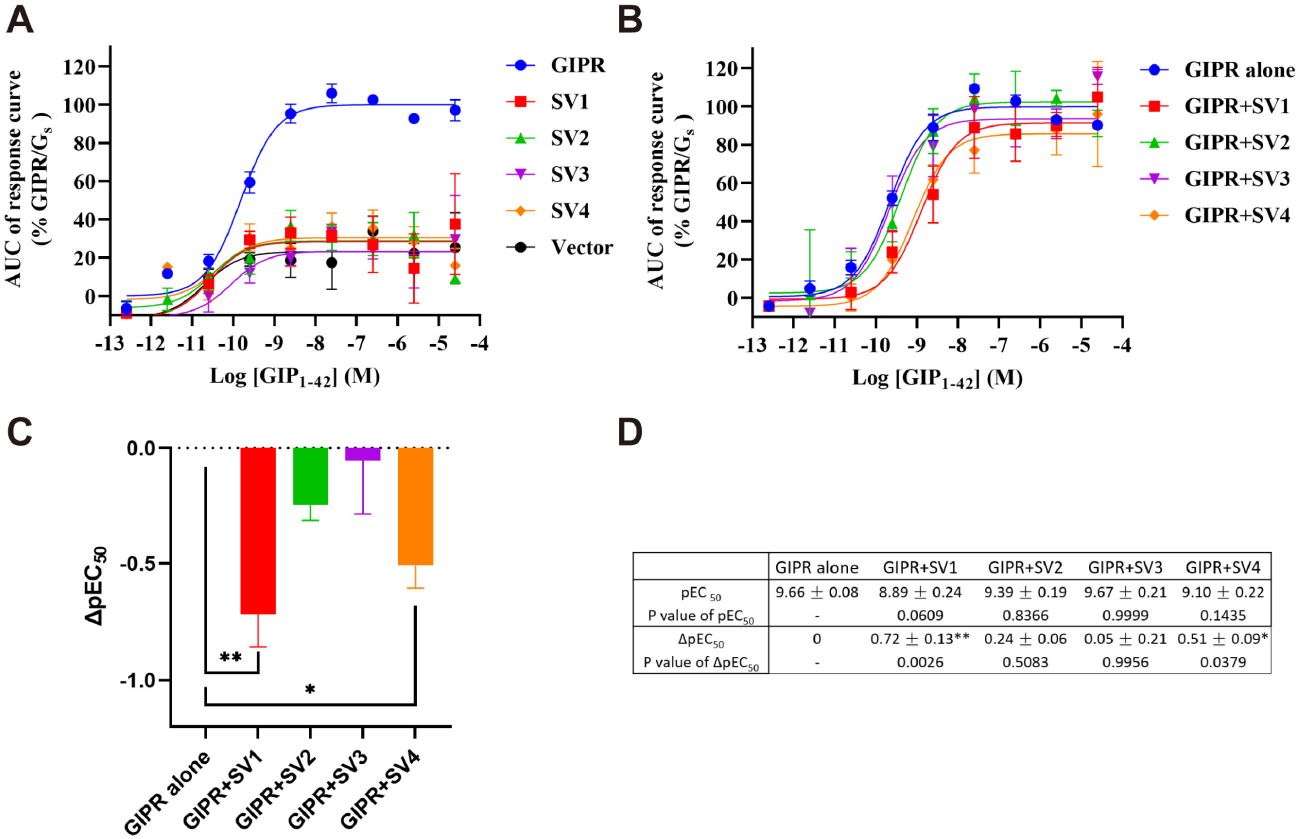
Effects of GIPR splice variants (SVs) on the wild-type (WT) GIPR mediated G_s_ protein coupling in HEK293T cells co-expressing GIPR and individual SVs. **A**, GIP_1-42_-induced G_s_ coupling of individually expressed SVs and GIPR. Concentration-response curves are expressed as area-under-the-curse (AUC) across the time-course response curve (0 to 13.5 min) for each concentration and normalized to WT GIPR. **B**, Effects of SVs on GIP_1-42_ induced G_s_ protein coupling at GIPR. **C**, EC_50_ differences of GIPR mediated G_s_ protein coupling under the influence of SVs. **D**, G_s_ protein coupling profiles of GIPR affected by SVs. Cells were co-transfected with GIPR and each SV at a 1:3 ratio. Data shown are means ± SEM of six independent experiments performed in duplicate. Signals were normalized to the maximum (max) response of GIPR and concentration-response curves were analyzed using a three-parameter logistic equation. *P < 0.05; **P < 0.01. The following figure supplements are available for figure 6: Source data 6. Effects of GIPR splice variants (SVs) on the wild-type (WT) GIPR mediated G_s_ protein coupling in HEK293T cells co-expressing GIPR and individual SVs.

### Diminished interaction between splice variants and signaling partners

As shown in Figure 7A-C, the helix 8 (H8) of SV1 adopted a distinct conformation from that of GIPR during molecular dynamics (MD) simulations, bent upwards and moved away from Gβ, thereby resulting in a reduced receptor-Gβ interface area. Of note, the specific residue in GIPR H8 (such as R405) stabilized the Gβ binding, which was absent in SV1, consistent with its role in G_s_-mediated signaling. Compared to GIPR, obvious differences in peptide-binding and β-arrestin 1 interface were observed for SV3 (Figure 7D-F). By replacing the GIPR extracellular domain (ECD) with a smaller domain (61 fewer residues), SV3 reorganized its extracellular half including ECL1 to accommodate peptide binding with a smaller peptide-receptor interface (from 3,280 Å^2^ in the last 500 ns MD simulation of GIPR to 2,826 Å^2^ in that of SV1). As far as the intracellular half is concerned, β-arrestin 1 inserted deeper to the SV3 core compared with that of GIPR (Figure 7E). These different structural and dynamics features between SV3 and GIPR highlight their distinct signaling properties.

**Figure 7.**
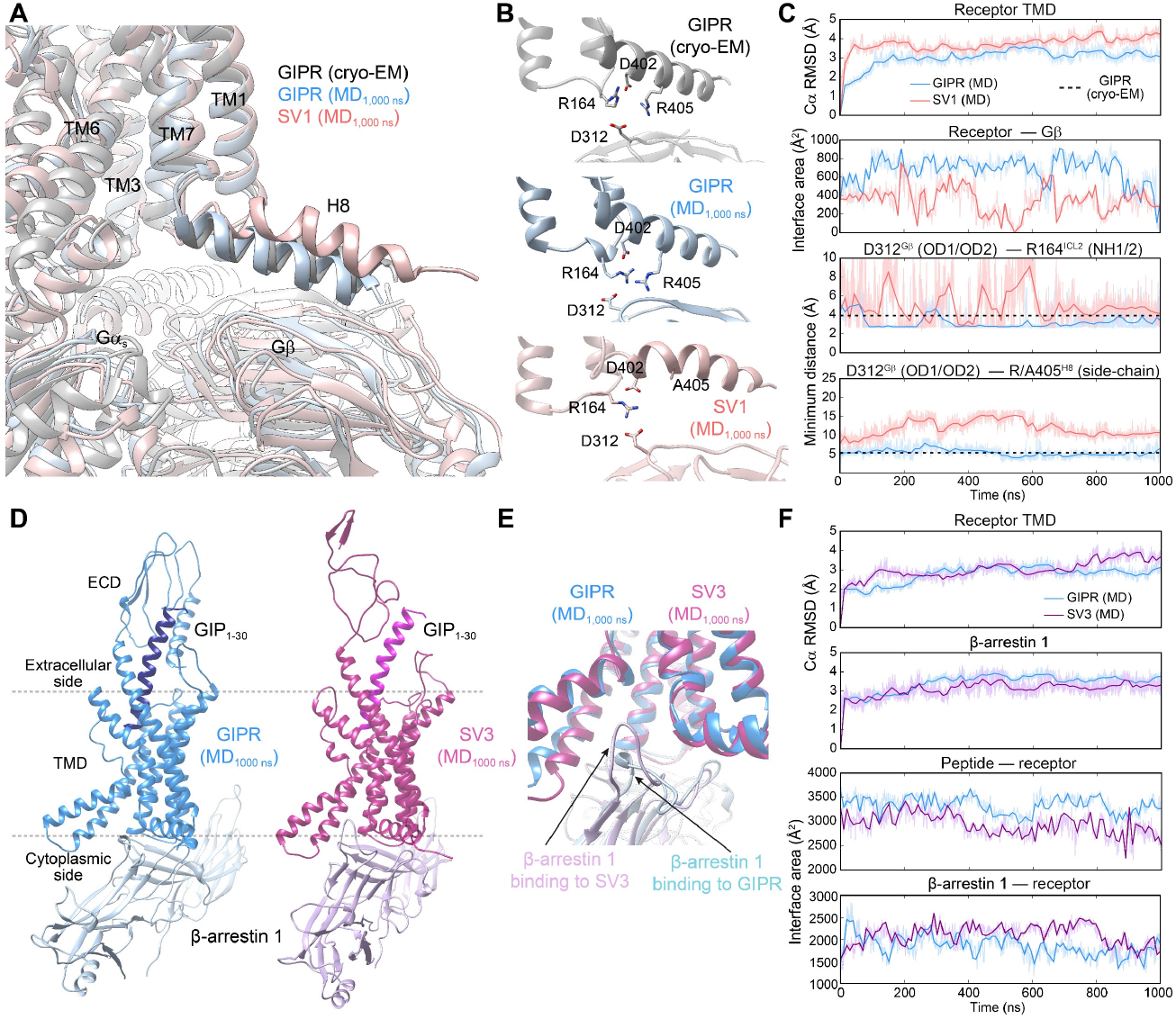
Molecular dynamics (MD) simulations of SV1 and SV3. **A**, Comparison of receptor-G_s_ conformation between the cryo-EM structure and final simulation snapshots of SV1 and GIPR. **B**, Comparison of the H8–Gβ interface between GIPR and SV1. **C**, Analysis of the MD simulation trajectories in (**A)**: top, root mean square deviation (RMSD) of Cα positions of the receptor TMD, where all snapshots were superimposed on the cryo-EM structure of both GIP- and G_s_-bound GIPR TMD (PDB code: 7DTY) using the Cα atom; upper middle, the buried surface area between receptor and Gβ, interface areas were calculated using freeSASA; lower middle, minimum distances between the charged atoms of D312^Gβ^ and R164^ICL2^ during MD simulations; bottom, minimum distances between the charged atoms of D312^Gβ^ and the side-chain atoms of R405^H8^ (GIPR) or A405^H8^ (SV1) during MD simulations. **D**, Comparison of the final MD snapshots between GIPR-β-arrestin 1 and SV3-β-arrestin 1. **E**, Different β-arrestin 1 modes between GIPR and SV3 according to MD simulations. **F**, Analysis of the MD simulation trajectories in (**D)**: top, Cα RMSD of the receptor TMD, where all snapshots were superimposed on the cryo-EM structure of both GIP- and G_s_-bound GIPR TMD (PDB code: 7DTY); upper middle, Cα RMSD calculation for β-arrestin 1, where all snapshots were superimposed on the cryo-EM structure of β_1_AR-bound β-arrestin 1 (PDB code: 6TKO) using the Cα atom; lower middle, the buried surface area between receptor and peptide during MD simulations; bottom, the buried surface area between receptor and β-arrestin 1 during MD simulations.

## Discussion

GIP, glucagon-like peptide-1 (GLP-1) and glucagon (GCG) together play a pivotal role in glucose homeostasis mediated via their respective receptors (Cho, Merchant, & Kieffer, 2012; Sekar, Singh, Arokiaraj, & Chow, 2016). GCG increases the release of glucose, while GIP and GLP-1R work synergistically to cause postprandial insulin secretion, regulate glucagon secretion, stimulate β cell proliferation and protect it from apoptosis (Alexiadou, Anyiam, & Tan, 2019; Estall & Drucker, 2006; Hansotia & Drucker, 2005; Seino, Fukushima, & Yabe, 2010; Skow, Bergmann, & Knop, 2016). Of note is that GIP promotes the release of both insulin and glucagon (Gasbjerg et al., 2018) thereby modulating the action of GLP-1 and GCG on sugar metabolism, probably involving some SVs of GIPR.

A common feature of the SVs examined is that they neither bind the native ligand, GIP_1-42_, nor elicit signal transduction. When co-expressed with WT GIPR, all of them reduced peptide binding in a similar manner while displaying distinct signaling profiles (Figure 8). Both SV1 and SV4 decreased GIPR-mediated cAMP and β-arrestin 2 responses; SV3 selectively suppressed **β**-arrestin 2 recruitment, and SV2 had no effect on the two signaling events, but diminished GIPR presence in the membrane. While SV1 (SV4 to less extent) may have the preference for activating the G_s_ pathway, SV3 obviously is biased towards β-arrestin 2 recruitment.

**Figure 8.**
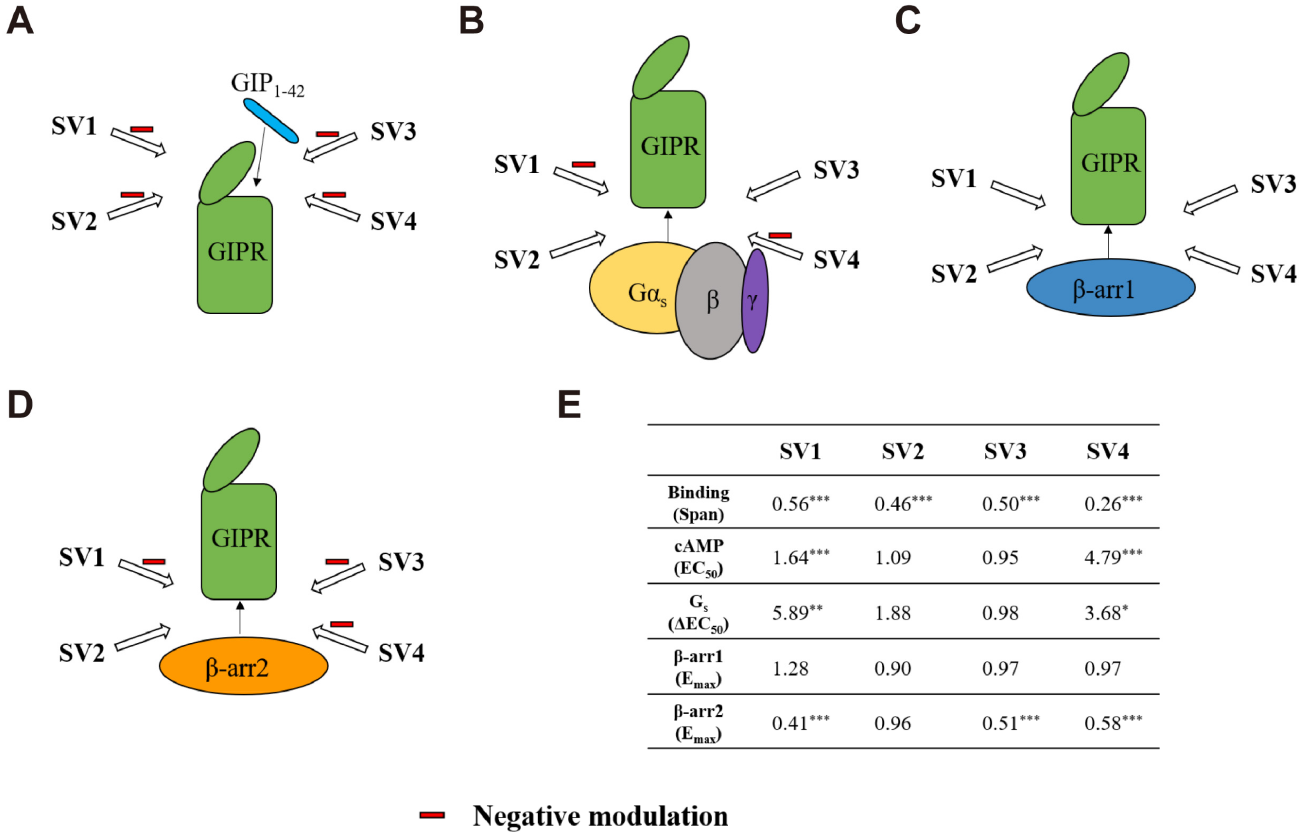
Characterization of the effects of SVs on ligand binding and signaling profiles in HEK293T cells co-expressing GIPR and individual SVs. **A-D**, Schematic diagram representing the effects of SVs on ligand binding (**A**), cAMP signaling and G_s_ activation (**B**), β-arrestins 1 (β-arr1) (**C**) and 2 (β-arr2) recruitment (**D**). **E**, Ratio of parameters of ligand binding and signaling to GIPR alone. One-way ANOVA were used to determine statistical difference (*P< 0.05, **P< 0.01, ***P< 0.0001).

Consistent with previous findings showing that SVs are capable of altering signaling profiles compared to WT receptors (Kochman, 2014; Maggio et al., 2016), our data suggest a constitutive biased mechanism different from signal bias caused by various ligands. For example, SVs of the C-X-C chemokine receptor 3 (CXCR3) could activate different signaling pathways through biased agonism (Berchiche & Sakmar, 2016), and SV1 of the growth hormone–releasing hormone receptor (GHRHR) preferentially transduces signals via β-arrestins while GHRHR predominantly activates G_s_ proteins (Cong et al., 2021). However, unlike other GPCR SVs, that of GIPR are incapacitated in terms of ligand-binding and signal transduction *per se*, but negatively affect that of the WT receptor in a ligand-independent and signaling biased manner.

Bidirectional regulation of carbohydrate levels by GIP_1-42_ is essential to the maintenance of glucose homeostasis, although this hinders the development of therapeutic agents targeting GIPR (Killion et al., 2018). It seems that such a unique modulation of gut hormone actions is finely tuned by SVs with differentiated functionalities: the repression of cAMP response imposed by RAMP3 could be strengthened by SV1 and SV4, whereas β-arrestin 2 signaling is solely modified by SV3. Unlike the other three SVs, SV2 appears as a sequester that redistributes the membrane GIPR to the cytoplasm, evidenced by immunofluorescence staining when both WT GIPR and SV2 are co-expressed. Whether this constitutes a shutdown mechanism for GIPR function remains to be explored.

The roles of SVs in GIPR functioning are unique not only because they are of repressive nature but also due to their synergistic actions with RAMP3 that itself is a negative regulator of most members of the glucagon receptor subfamily of class B1 GPCRs (Shao et al., 2021). It remains elusive if the above described phenomenon constitutes a “doubly insured” mechanism for signal modulation in order to fine tune the action of GIP_1-42_. Clearly both in-depth structural and biochemical studies are required to solve the puzzle.

## Materials and Methods

### Key resource table

| **Reagent type**<br>**(species) or**<br>**resource** | **Designation** | **Source or reference** | **Identifier** | **Additional**<br>**information** |
| --- | --- | --- | --- | --- |
| Transcript ID | WT GIPR | ENSEMBL | ENST00000590918.6 |  |
| Transcript ID | Splice variant 1 | ENSEMBL | ENST00000263281.7 |  |
| Transcript ID | Splice variant 2 | ENSEMBL | ENST00000304207.12 |  |
| Transcript ID | Splice variant 3 | ENSEMBL | ENST00000652180.1 |  |
| Transcript ID | Splice variant 4 | ENSEMBL | ENST00000585889.1 |  |
| Cell line (*Homo sapiens*) | HEK293T | ATCC | Cat. No.: CRL-3216 |  |
| Cell line (hamster) | CHO-K1 | ATCC | Cat. No.: CCL-61 |  |
| Recombinant DNA reagent | pcDNA3.1-GIPR (WT and splice variants) | This paper | N/A |  |
| Recombinant DNA reagent | pcDNA3.1-GIPR (WT)-HA | doi: 10.1016/j.bcp.2020.114001 | N/A |  |
| Recombinant DNA reagent | pcDNA3.1-GIPR (splice variants) -FLAG | This paper | N/A |  |
| Recombinant DNA reagent | pcDNA3.1-GIPR (WT and splice variants)- | This paper | N/A |  |
|  | Rluc8 (Renilla luciferase 8) |  |  |  |
| Recombinant DNA reagent | Venus-β-arrestins 1 and 2 | doi:<br>10.1016/j.bcp.2020.114001 | N/A | |
| Recombinant DNA reagent | pcDNA3-RAMP3 | doi:<br>10.1016/j.apsb.2021.07.028 | N/A |  |
| Peptide, recombinant protein | GIP<sub>1-42</sub> | GenScript | N/A |  |
| Chemical compound, drug | Bovine serum albumin (BSA) | ABCONE | Cat. No.:<br>A23088-100G |  |
| Chemical compound, drug | 3-Isobutylene-1-methylxanthine (IBMX) | ABCONE | Cat. No.:<br>I72182-250MG |  |
| Chemical compound, drug | Fetal bovine serum (FBS) | Gibco | Cat. No.:<br>10099-141 |  |
| Chemical compound, drug | Dulbecco's modified Eagle's medium (DMEM) | Gibco | Cat. No.:<br>12430-054 |  |
| Chemical compound, drug | Ham's F-12 nutrient mix (F12) | Gibco | Cat. No.:<br>C11765500 |  |
| Chemical compound, drug | Hanks' Balanced Salt solution (HBSS) | Gibco | Cat. No.:<br>C14175500 |  |
| Chemical compound, drug | HEPES | Gibco | Cat. No.:<br>15630-080 |  |
| Chemical compound, drug | Sodium pyruvate | Gibco | Cat. No.:<br>11360-0'70 |  |
| Chemical compound, drug | Lipofectamine 2000 transfection reagent | Invitrogen | Cat. No.:<br>11668-019 |  |
| Chemical compound, drug | <sup>125</sup>I-GIP | PerkinElmer | Cat. No.:<br>NEX40201<br>0UC |  |
| Chemical compound, drug | Coelenterazine H | Yeasen Biotech | Cat. No.:<br>40906ES02 |  |
| Antibody | Anti-FLAG primary antibody | Sigma-Aldrich | Cat. No.:<br>F3165 |  |
| Antibody | Anti-HA primary antibody | Cell Signaling Technology | Cat. No.:<br>3724S |  |
| Antibody | Anti-mouse Alexa Fluor 647 conjugated secondary antibody | Invitrogen | Cat. No.:<br>A31571 |  |
| Antibody | Anti-rabbit Alexa Fluor 488 conjugated secondary antibody | Invitrogen | Cat. No.:<br>A21206 |  |
| Commercial assay, kit | LANCE Ultra cAMP kit | PerkinElmer | Cat. No.:<br>2675984 |  |
| Software, algorithm | GraphPad Prism v8.4 | GraphPad Software | N/A | https://www.graphpad.com/ |
| Software, algorithm | FreeSASA | doi:<br>10.12688/f1000research.7931.1 | N/A | http://freesasa.github.io/ |
| Software, algorithm | Gromacs 2020.1 | doi:<br>10.1016/j.softx.2015.06.001 | N/A | https://manual.gromacs.org/2020.1/download.html |
| Software,<br>algorithm | Protein Preparation Wizard | Schrödinger | N/A | https://www.schrodinger.com/products/protein-preparation-wizard |
| Software,<br>algorithm | CHARMM-GUI Membrane Builder v3.6 | doi:<br>10.1002/jcc.23702 | N/A | https://charmm-gui.org/ |
| Software,<br>algorithm | CHARMM36-CAMP | doi:<br>10.1021/ct200328p | N/A |  |
| Software,<br>algorithm | LINCS algorithm | doi:<br>10.1021/ct700200b | N/A |  |
| Software,<br>algorithm | Semi-isotropic Parrinello-Rahman barostat | doi: 10.1016/0022-3093(93)90111-A | N/A |  |

### Cell culture

HEK293T cells were maintained in DMEM (Gibco) supplemented with 10% FBS (Gibco) and 100 mM sodium pyruvate (Gibco). CHO-K1 cells were maintained in F12 (Gibco) supplemented with 10% FBS. All cells were incubated in a humidified environment at 37 **°** C in 5% CO_2_.

### Construct

cDNAs were inserted into pcDNA3.1 vector by one-step cloning. Addition of FLAG- and HA-tags to WT GIPR or SVs was carried out by site-directed mutagenesis. WT GIPR or SVs were cloned to the backbone of Rluc8 at the C terminus. All constructs were confirmed by DNA sequencing (GENEWIZ, Suzhou, China). To optimizing the co-transfection assays, three GIPR *vs*. SV ratios (1:1, 1:3 and 1:6) were tried. Since the impact of 1:1 on GIPR activity was hard to observe and that of 1:3 and 1.6 was similar, we selected 1:3 for the entire study.

### cAMP accumulation assay

GIP_1-42_ stimulated cAMP accumulation was measured by a LANCE Ultra cAMP kit (PerkinElmer). Cells were seeded onto 6-well cell culture plates and transiently transfected with 4 μg plasmid using Lipofectamine 2000 transfection reagent (Invitrogen). After 24 h culture, the transfected cells were seeded onto 384-well microtiter plates at a density of 3,000 cells per well in HBSS (Gibco) supplemented with 5 mM HEPES (Gibco), 0.1% (w/v) bovine serum albumin (BSA) and 0.5 mM IBMX (Sigma-Aldrich). The cells were stimulated with different concentrations of GIP_1-42_ for 40 min at room temperature (RT). Eu and Ulight were then diluted by cAMP detection buffer and added to the plates separately to terminate the reaction. Plates were incubated at RT for 40 min and the fluorescence intensity measured at 620 nm and 650 nm by an EnVision multilabel plate reader (PerkinElmer).

### Whole-cell binding assay

CHO-K1 cells were seeded at a density of 30,000 cells/well in Isoplate-96 plates (PerkinElmer). The WT GIPR or SVs were transiently transfected using Lipofectamine 2000 transfection reagent. Twenty-four hours after transfection, cells were washed twice, and incubated with blocking buffer (F12 supplemented with 33 mM HEPES and 0.1% BSA, pH 7.4) for 2 h at 37°C. For homogeneous binding, cells were incubated in binding buffer with a constant concentration of ^125^I-GIP (40 pM, PerkinElmer) and increasing concentrations of unlabeled GIP_1-42_ (3.57 pM to 1 μM) at RT for 3 h. Following incubation, cells were washed three times with ice-cold PBS and lysed by addition of 50 μL lysis buffer (PBS supplemented with 20 mM Tris-HCl, 1% Triton X-100, pH 7.4). Fifty μL of scintillation cocktail (OptiPhase SuperMix, PerkinElmer) was added and the plates were subsequently counted for radioactivity (counts per minute, CPM) in a scintillation counter (MicroBeta2 Plate Counter, PerkinElmer).

### β-arrestin 1/2 recruitment

HEK293T cells (3.5×10^6^ cells/mL) were grown for 24 h before transfection with 4 μg plasmid containing a GIPR/SV-Rluc8:Venus-β-arrestin1/2 at ratio of 1:9, or a GIPR-Rluc8:SV:Venus-β-arrestin1/2 at a ratio of 1:3:9. Transiently transfected cells were then seeded onto poly-D-lysine coated 96-well culture plates (50,000 cells/well) in DMEM with 10% FBS. Cells were grown overnight before incubation in assay buffer (HBSS supplemented with 10 mM HEPES and 0.1% BSA, pH 7.4) for 30 min at 37°C. Coelentrazine-h (Yeasen Biotech) was added to a final concentration of 5 μM for 5 min before bioluminescence resonance energy transfer (BRET) readings were made using an EnVision plate reader. BRET baseline measurements were collected for 15 cycles prior to ligand addition. Following peptide addition, BRET was measured for 55 cycles. The BRET signal (ratio of 535 nm over 470 nm emission) was corrected to the baseline and then vehicle-treated condition to determine ligand-induced changes in BRET response. Concentration-response values were obtained from the area-under-the-curve (AUC) of the responses elicited by GIP_1-42_.

### Immunofluorescence staining

HEK293T cells were seeded in 6-well plates and transfected with 4 μg plasmid containing GIPR-HA or/and SV-FLAG. After 24 h, cells were collected and reseeded in 96-well plates until they reached 50%∼70% confluence. Cells were washed with PBS before fixation with 4% paraformaldehyde for 15 min. Then they were washed three more times and blocked with 5% BSA plus 0.1% Triton X-100 for 1 h at RT. Rabbit anti-HA primary antibody (diluted 1:500) or/and mouse anti-FLAG primary antibody (diluted 1:300) were diluted with incubation buffer (PBS supplemented with 5% BSA) for 1 h followed by 3-time wash. Cells were reacted with 200 μL interaction buffer containing donkey anti-rabbit Alexa 488-conjugated secondary antibody or/and donkey anti-mouse Alexa 647-conjugated secondary antibody (diluted 1:1000) at RT for 1 h in the dark. After final washing, nuclei were stained with Hoechst 33258 for 5 min. Cells were imaged using a high-resolution microscope DeltaVision™ Ultra (GE Healthcare, Boston, USA).

### G protein NanoBiT assay

HEK293T cells (3.5×10^6^ cells/mL) were grown for 24 h to reach 70% to 80% confluence. Then the cells were transiently transfected with GIPR, Gα_s_-LgBiT, Gβ1, and Gγ2-SmBiT at a 2:1:5:5 mass ratio, or GIPR, SV, Gα_s_-LgBiT, Gβ1, and Gγ2-SmBiT at a 2:6:1:5:5 mass ratio. Twenty-four hours after transfection, cells were seeded into poly-D-lysine coated 96-well culture plates at a density of 50,000 cells per well in DMEM with 10% FBS. Cells were grown overnight before incubation in HBSS buffer (pH 7.4) supplemented with 0.1% BSA and 10 mM HEPES for 30 mins at 37°C (no CO_2_). They were then reacted with coelenterazine H (5 μM) for 2 h at RT. Luminescence signals were measured using an EnVision plate reader at 15-s intervals (25°C). Briefly, following the baseline reading for 3.5 min, GIP_1-42_ was added, and the reading continued for 13.5 min. Data were corrected to baseline and vehicle-treated samples. The area-under-the-curve (AUC) across the time-course response curve was determined and normalized to the WT GIPR, which was set to 100%.

### Molecular dynamics simulation

Molecular dynamic simulations were performed by Gromacs 2020.1. The homology models of SV1 and SV3 were generated using the cryo-EM structure of the full-length GIPR (PDB code: 7DTY) (Zhao et al., 2021) and the X-ray structure of the GIPR ECD (PDB code: 2QKH) (Parthier et al., 2007) as templates. All peptide–receptor–G_s_ complexes were built based on the cryo-EM structure of the GIP–GIPR–G_s_ complex (PDB code: 7DTY) (Zhao et al., 2021) and prepared by the Protein Preparation Wizard (Schrodinger 2017-4) with the Nb35 nanobody removed. To build the model of peptide–receptor–β-arrestin 1 complex, the receptor in complex with both peptide and G_s_ was aligned to the published β-arrestin 1–bound β_1_AR structure (PDB code: 6TKO) (Lee et al., 2020). The receptor chain termini were capped with acetyl and methylamide. The residues G2 and C3 of Gα_s_ were *N-*myristoylated and palmitoylated, respectively (Kato et al., 2019). All missing backbone and side chains were modeled using Prime (Schrodinger 2017-4) and the titratable residues were left in their dominant state at pH 7.0. To build MD simulation systems, the complexes were embedded in a bilayer composed of 254∼315 POPC lipids and solvated with 0.15 M NaCl in explicit TIP3P waters using CHARMM-GUI Membrane Builder v3.6 (Wu et al., 2014). The CHARMM36-CAMP force filed (Guvench et al., 2011) was adopted for protein, peptides, lipids and salt ions. The Particle Mesh Ewald (PME) method was used to treat all electrostatic interactions beyond a cut-off of 12 Å and the bonds involving hydrogen atoms were constrained using LINCS algorithm (Hess, 2008). The complex system was first relaxed using the steepest descent energy minimization, followed by slow heating of the system to 310 K with restraints. The restraints were reduced gradually over 50 ns. Finally, restrain-free production run was carried out for each simulation, with a time step of 2 fs in the NPT ensemble at 310 K and 1 bar using the Nose-Hoover thermostat and the semi-isotropic Parrinello-Rahman barostat (Aoki & Yonezawa, 1992), respectively. The buried interface areas were calculated with FreeSASA using the Sharke-Rupley algorithm with a probe radius of 1.2 Å (Mitternacht, 2016).

### Statistical analysis

Statistical analysis was performed using GraphPad Prism 8.4 (GraphPad Software). For signaling assays, data of individual experiments were normalized to the maximum responses in cells expressing only the WT GIPR. Non-linear curve fit was performed using a three-parameter logistic equation (log (agonist *vs*. response)). All data are presented as means ± SEM. Significant differences were determined by one-way ANOVA with Dunnett’s test. For co-localization analysis, Pearson’s correlation coefficients (r) were performed using the co-localization threshold plugin of ImageJ. Five separate Regions of Interest (ROI) were selected and means ± SEM were determined.

## Acknowledgments

We are grateful to Dr. Zhaotong Cong, Ms. Yan Chen and Mr. Chao Zhang for technical assistance. This work was partially supported by National Natural Science Foundation of China 81872915 (M.-W.W.), 82073904 (M.-W.W.), 82121005 (D.Y.), 81973373 (D.Y.) and 21704064 (Q.Z.); National Science & Technology Major Project of China– Key New Drug Creation and Manufacturing Program 2018ZX09735–001 (M.-W.W.) and 2018ZX09711002–002–005 (D.Y.); National Science & Technology Major Project of China – Innovation 2030 for Brain Science and Brain-Inspired Technology 2021ZD0203400 (Q.Z.); the National Key Basic Research Program of China 2018YFA0507000 (M.-W.W.); Novo Nordisk-CAS Research Fund grant NNCAS-2017–1-CC (D.Y.) and SA-SIBS Scholarship Program (D.Y.).

## Author contributions

D.Y. and M.-W.W. designed research; K.H., L.S., F.Z., A.D., and X.C. performed research; K.H., L.S., Q.Z., R.C.S. and D.Y. analyzed data; and K.H., L.S., and M.-W.W. wrote the paper.

## Data availability

All data generated or analyzed during this study are included in the manuscript. Source data files have been provided for Figures 2-6.

